# Spatiotemporal proteomic profiling of genome editing-associated microenvironment

**DOI:** 10.64898/2026.10.23.760596

**Authors:** Siqi Xi, Linghao Kong, Weijie Zhou, Xuan Liang, Peng R. Chen, Xinyuan Fan

## Abstract

Genome editing outcomes are influenced by local genomic context, yet the dynamic protein assemblies that form around genome editing sites remain poorly characterized. Here, we developed a Cas9-TurboID proximity labeling platform that combines programmable genomic targeting with rapid and minimally perturbative protein labeling, enabling spatiotemporal mapping of dynamic protein assemblies associated with distinct genome editing processes. Dynamic comparative profiling identified ACTL6A and H2A.Z as candidate regulators of genome editing, and Cas9 fusion with either ACTL6A or H2A.Z enhanced editing efficiency and improved editing outcomes in heterochromatic regions, which are generally refractory to genome editing due to limited DNA accessibility. We further extended the platform to prime editing and identified WRN as a functional regulator. Together, this study provides a strategy for systematically decoding editor-associated protein microenvironments, establishing a framework for understanding genome editing mechanisms and guiding the rational engineering of genome editors.

---

Genome editing technologies based on CRISPR-Cas systems have revolutionized genome engineering and are increasingly being applied in therapeutic applications^1,2^. Although newer platforms such as base editing (BE) and prime editing (PE) improve editing precision by avoiding or minimizing double-strand breaks, editing efficiency and repair outcomes remain influenced by the local genomic context^3,4^. Previous studies have demonstrated that editing efficiency and DNA repair outcomes are influenced by chromatin accessibility, epigenetic state, DNA sequence composition, and other locus-specific features^5–11^. Together, these studies suggest that the local chromatin environment plays a central role in regulating DNA repair during genome editing.

Despite extensive characterization of genomic and epigenomic determinants, the molecular mechanisms by which local genomic context influences genome editing remain incompletely understood. These contextual features ultimately exert their effects through the dynamic recruitment and organization of proteins at DNA lesions, including chromatin remodelers, histone regulators, and DNA repair factors. The composition and temporal dynamics of these local protein assemblies are likely to influence repair pathway choice and editing outcomes^12,13^. However, how these protein microenvironments are established and remodeled during genome editing remains largely unknown, particularly across editing modalities that generate distinct DNA repair intermediates, including Cas9-induced double-strand breaks (DSBs), nickase-induced single-strand breaks (SSBs), and prime editing.

Recent approaches have combined catalytically inactive Cas9 (dCas9) with proximity-labeling enzymes to profile proteins associated with predefined genomic loci, providing valuable insights into locus-specific chromatin composition^14^. However, these approaches primarily interrogate static genomic locations and therefore cannot capture the dynamic remodeling of local protein environments that accompanies active DNA cleavage and repair. Such remodeling is expected to differ substantially among genome-editing modalities that generate distinct DNA intermediates, including Cas9-induced DSBs, nickase-induced SSBs, and prime editing. A systematic characterization of these local proteomic landscapes is therefore needed to elucidate DNA repair mechanisms during genome editing and to identify opportunities for improving editing efficiency and precision.

Proximity labeling technologies, such as BioID and APEX, have greatly expanded the ability to profile the local protein environment. However, resolving the dynamic protein assemblies formed during genome editing requires labeling approaches that balance efficiency, temporal resolution, and minimal perturbation^15–17^. Here, we employed the proximity labeling enzyme TurboID^18^, which enables non-invasive proximity labeling with minimal perturbation of endogenous DNA damage responses, fused to Cas9 to achieve locus-specific DNA repair microenvironment labeling. Using Cas9 and Cas9(D10A) to induce site-specific DSBs and SSBs, respectively, we characterized the temporal dynamics of proteins recruited around DNA lesions at both the exogenous reporter locus and endogenous genomic target. Two proteins, ACTL6A and H2A.Z, were found to improve Cas9 editing, especially in heterochromatin. Finally, we extended this approach to prime editing, uncovering the recruitment of chromatin remodelers and DNA secondary-structure-binding proteins associated with complex editing intermediates, and identified WRN as a potential endogenous regulator of prime editing. Taken together, we established a strategy for profiling the dynamic protein microenvironments associated with genome editors, enabling systematic characterization of the proteins that influence genome editing outcomes.

## Results

### Cas9-TurboID assay for DSB and SSB repair microenvironment labeling

To map the dynamic protein microenvironment during target site repair, we directly fused wild-type Cas9 or the nickase variant Cas9(D10A) with TurboID to construct proximity-labeling tools capable of generating double-strand breaks (DSBs) and single-strand breaks (SSBs), respectively. The corresponding repair pathways are illustrated (**Fig. 1a**). We first evaluated their cleavage activity and repair kinetics using firefly luciferase-based reporters (fLuc) designed to represent DSB and SSB repair outcomes (**Fig. 1b)**^19^. The time-resolved bioluminescence measurements, combined with indel frequency analysis, revealed the repair kinetics following DSB or SSB induction. Repair-associated bioluminescence signals approached saturation by 48 h, whereas the indel frequencies continued to accumulate throughout the time course, suggesting that repeated cycles of Cas9 cleavage and repair persist after the initial repair response (**Fig. 1c-d)**.

**Fig. 1.**
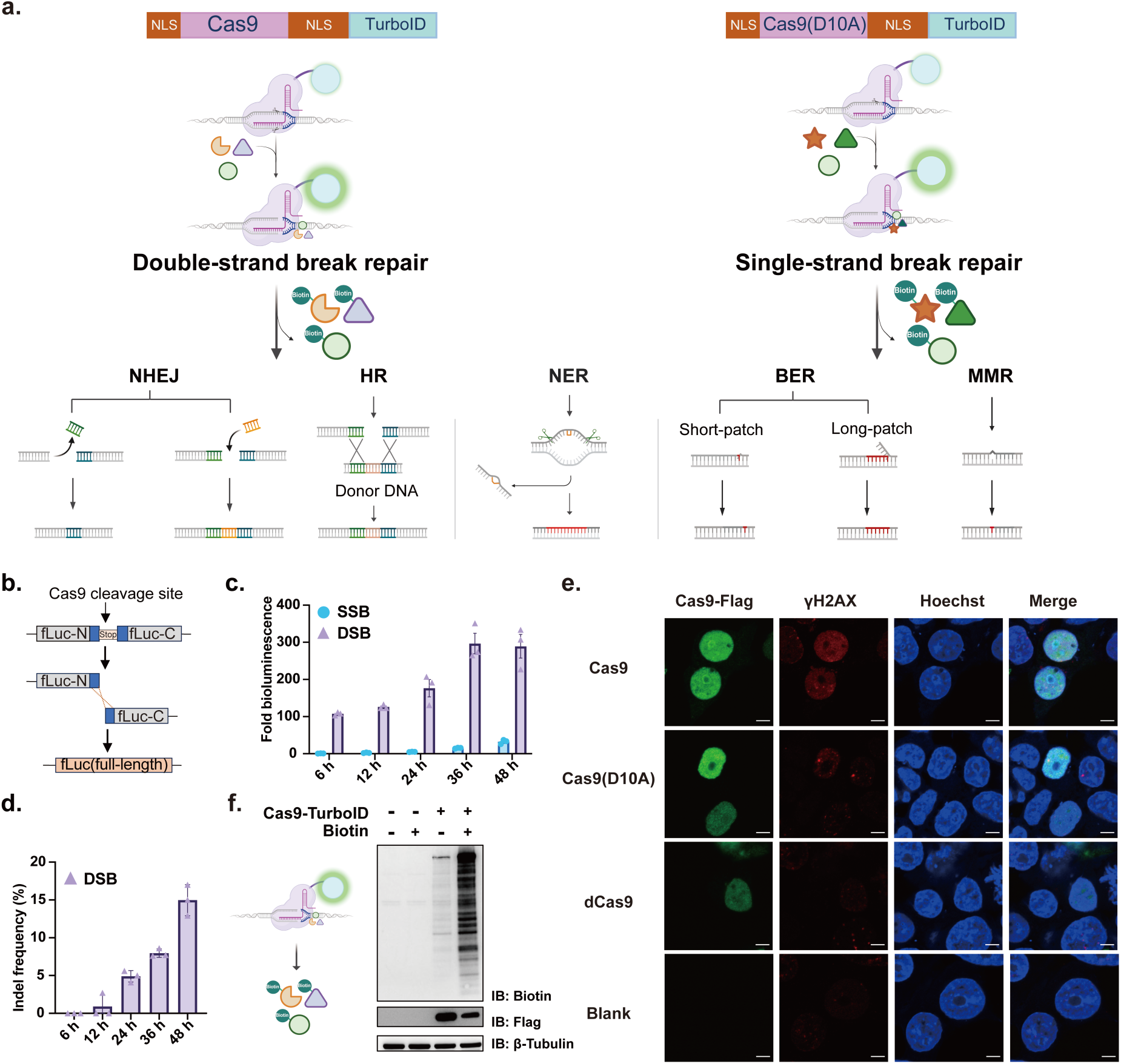
Cas9-TurboID assay for DSB and SSB repair microenvironment labeling. **a,** Schematic of DNA repair pathways following genome editing by Cas9-TurboID or Cas9(D10A)-TurboID. Cas9-TurboID induces double-strand breaks (DSBs) repaired primarily through non-homologous end joining (NHEJ) and homology-directed repair (HR), whereas Cas9(D10A)-TurboID generates single-strand breaks (SSBs) repaired via base excision repair (BER) and mismatch repair (MMR). An additional repair pathway, nucleotide excision repair (NER), was also shown. Created with BioRender.com. **b,** Schematic of firefly luciferase-based reporter (fLuc) systems used to quantify DSB and SSB repair efficiency. **c-d,** Time-resolved bioluminescence signals following DSB and SSB induction and indel frequencies of DSB induction at the Luciferase reporter site (*Luc*). Bioluminescence reflects repair reporter activity, while indel frequencies quantify cumulative editing outcomes. Data are shown as mean ± s.d. (n = 3 biological replicates). **e,** Immunofluorescence analysis of γH2AX foci in HEK293T cells expressing Cas9, Cas9(D10A), or dCas9, with untransfected cells as the blank control. Scale bar, 5 µm. Anti-rabbit-Alexa Fluor 488 for Cas9-Flag and anti-mouse-Alexa Fluor 555 for γH2AX. **f,** Immunoblotting analysis of biotinylated proteins demonstrating efficient TurboID-mediated proximity labeling in Cas9-TurboID-expressing cells. Cas9-TurboID-Flag was used for labeling, and β-Tubulin was used as a loading control.

Immunofluorescence analysis of γH2AX foci showed that the DNA damage occurred in wild-type Cas9 (Cas9-WT)^20^. Unexpectedly, increased γH2AX signals were also observed under the Cas9(D10A) condition, although to a lesser extent than in the Cas9-WT condition, suggesting that nick-induced lesions can also trigger DNA damage responses, potentially through unintended secondary cleavage events^21^. In contrast, γH2AX levels in dCas9-expressing cells remained equivalent to basal background levels, indicating that Cas9 occupancy alone does not induce substantial DNA damage signaling (**Fig. 1e**). Furthermore, immunoblotting analysis verified efficient, biotin-dependent proximity labeling by Cas9-TurboID (**Fig. 1f**). Having confirmed that the Cas9-TurboID fusion constructs retain high editing efficiency and robust proximity-labeling capacity, we applied the Cas9-TurboID for subsequent time-resolved proteomic experiments.

### Dynamic proteomic profiling of DSB and SSB repair microenvironment at exogenous reporter locus

To dissect the dynamic protein recruitment during the repair process, cells expressing Cas9-TurboID or Cas9(D10A)-TurboID were subjected to biotin labeling at the indicated time, followed by proteomic analysis of proximity-labeled proteins at the firefly luciferase-based reporter assay (Luc assay) (**Fig. 2a**). Comparative volcano plots with indicated pathway-related proteins revealed distinct protein enrichment kinetics between DSB and SSB conditions (**Fig. 2b**). The pathway proteins list was compiled from the Reactome database and REPAIRtoire^22,23^. Quantitative pathway analysis demonstrated an enrichment of SSB-associated repair factors at 12 h, whereas DSB-related factors peaked around 24 h, suggesting distinct temporal patterns of protein recruitment in SSB and DSB repair (**Fig. 2c**)^21,24,25^.

**Fig. 2.**
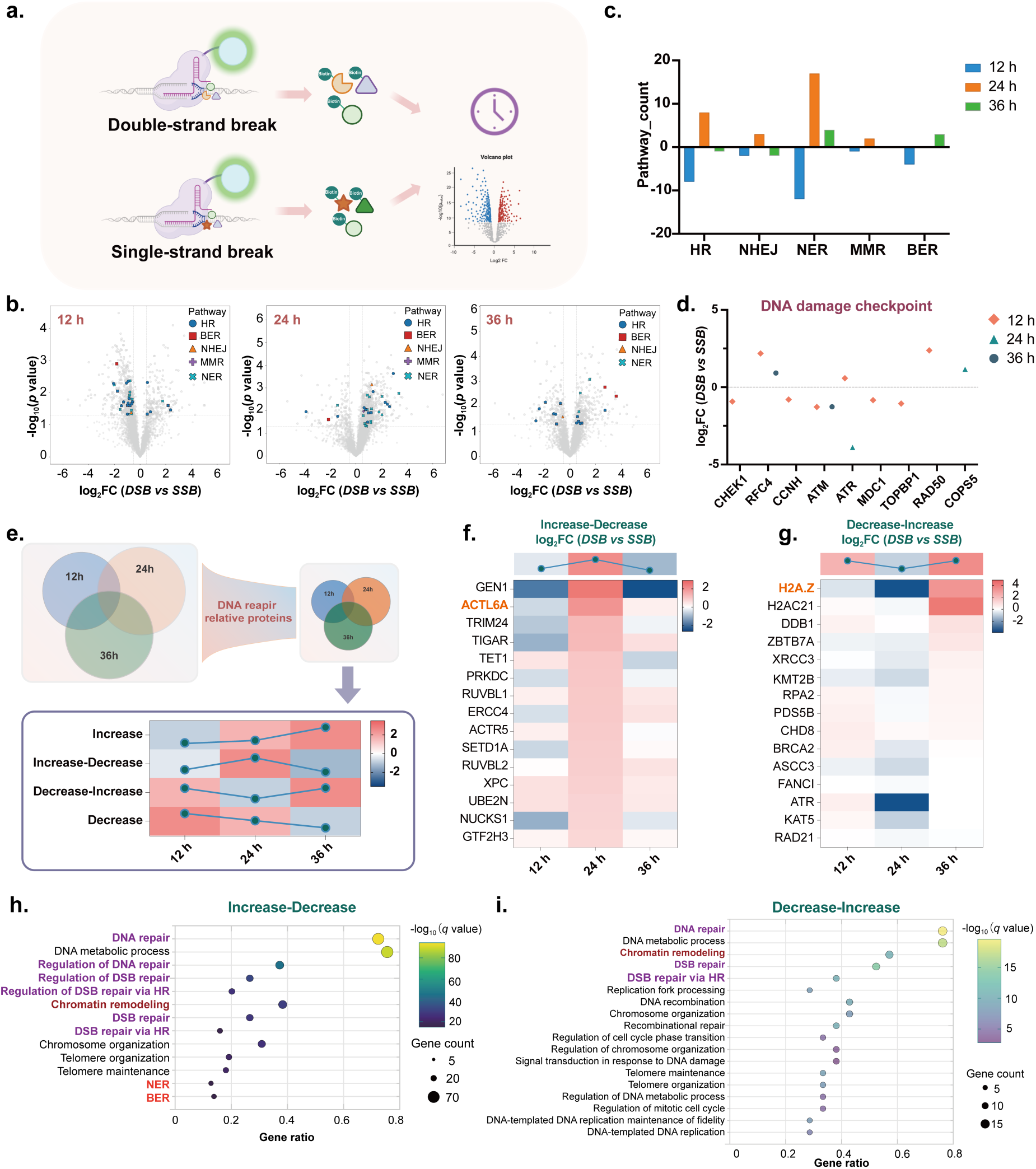
Proteomic profiling of DSB and SSB repair microenvironments at exogenous reporter locus. **a,** Schematic of the proximity-labeling strategy for capturing protein microenvironments associated with DSB and SSB repair for proteomic analysis at the firefly luciferase-based reporter assay (Luc assay). **b,** Volcano plots comparing protein enrichment profiles between DSB and SSB conditions at 12 h, 24 h, and 36 h. The pathway protein list is generated from the Reactome database. Differentially significantly enriched proteins were defined using *p* < 0.05 and |log_2_ Fold Change (FC)| ≥ 0.5. **c,** Summary of pathway-associated proteins identified from the volcano plots, showing their distribution and counts across 12 h, 24 h, and 36 h conditions. For each pathway, proteins with positive log₂FC (DSB *vs* SSB) values were classified as DSB-associated, whereas proteins with negative log₂FC values were classified as SSB-associated. **d,** Enriched DNA damage checkpoint proteins were analyzed at 12 h, 24 h, and 36 h, showing that most DNA damage checkpoint proteins were enriched at 12 h. Data are presented as the average value of log_2_FC (DSB *vs* SSB) at indicated time points from three independent biological replicates. **e,** Scheme for trend analysis. Proteins identified across all time points (12 h, 24 h, and 36 h) were first filtered based on DNA repair-related functions and subsequently subjected to trend analysis. The resulting trends were categorized into Increase, Increase-Decrease, Decrease-Increase, and Decrease patterns. **f-i,** Representative DNA repair-related proteins with different trends displaying distinct enrichment patterns between DSB and SSB conditions, together with corresponding Gene Ontology (GO) analysis^36^. **f** and **h** represent Increase-Decrease protein analysis; **g** and **i** represent Decrease-Increase protein analysis. Data are presented as the average value of log_2_ fold change of DSB *vs* SSB at 12 h, 24 h, and 36 h from three independent biological replicates.

Detailed temporal tracking revealed that the DNA damage checkpoint proteins exhibited preferential early recruitment at 12 h, consistent with an early-stage damage sensing response (**Fig. 2d**)^26,27^. This temporal pattern was followed by delayed recruitment of downstream repair factors: EXO1, which executes 5’ to 3’ end resection during homologous recombination (HR)^28,29^, and PRKDC, a core subunit of the non-homologous end joining (NHEJ) complex, both exhibited maximal enrichment at 24 h (**Extended Data Fig. 1a)**^30–32^. At 12 h, PARP2, an SSB-sensing protein and component of the SSB-repair assemblies^33^, was preferentially enriched under the SSB condition, consistent with its role in base excision repair (BER)^34^. Similarly, PNKP, a DNA end-processing enzyme, was preferentially enriched under the SSB condition at 12 h (**Extended Data Fig. 1b)**^35^. These early enrichments suggest that Cas9-induced SSB triggers a rapid and localized PARP-dependent repair response, consistent with canonical BER repair pathways. Additionally, nucleotide excision repair (NER) related proteins, which recognize DNA structure distortions, displayed enrichment at 12 h and 24 h (**Extended Data Fig. 1b**).

To systematically characterize the temporal dynamics of repair-associated proteins, we classified the dynamic recruitment profiles into four distinct temporal patterns: Increase, Increase-Decrease, Decrease-Increase, and Decrease (**Fig. 2e)**. Proteins exhibiting the Increase-Decrease pattern were enriched in BER and NER pathways according to GO analysis performed using Metascape^36^, consistent with the early engagement of SSB repair processes after nCas9 cleavage (**Fig. 2f**, **2h)**. This pattern also showed enrichment of DSB repair-associated pathways, including HR (**Fig. 2f**, **2h).** In contrast, proteins displaying the Decrease-Increase pattern showed preferential enrichment of DSB repair pathways (**Fig. 2g**, **2i**). Both patterns were associated with enrichment of chromatin remodeling-related proteins (**Extended Data Fig. 2**), highlighting the critical requirement of nucleosome accessibility during DNA repair. To determine whether Cas9 retention influences this local environment, we checked the Cas9 position after editing using CUT&RUN-qPCR. Compared to catalytically inactive dCas9, both Cas9 and Cas9(D10A) showed reduced target-site occupancy (**Extended Data Fig. 4a**), indicating a rapid dissociation and protein turnover at the target locus after editing.

### Spatiotemporal proteomic profiling reveals chromatin-remodeling constraints at endogenous genomic locus

To extend our findings to endogenous genomic contexts, we profiled DNA repair microenvironments at the canonical *EMX1* locus using Cas9-TurboID or Cas9(D10A)-TurboID, followed by temporal proximity labeling at 12 h, 24 h, and 36 h after editing initiation (**Fig. 3a**). Editing efficiency at the *EMX1* site was confirmed by indel frequency analysis (**Extended Data Fig. 3a**). Comparative volcano plots at 24 h showed a substantially stronger enrichment of proteins under the DSB condition relative to SSB (**Fig. 3b**). Pathway counting revealed reduced enrichment of pathway-related proteins at the endogenous locus compared to the exogenous Luc assay (**Fig. 3c**), reflecting the chromatin accessibility constraints imposed by native nucleosomal organization. Notably, pathway-associated protein recruitment patterns differ between endogenous and exogenous repair contexts (**Extended Data Fig. 3b**).

**Fig. 3.**
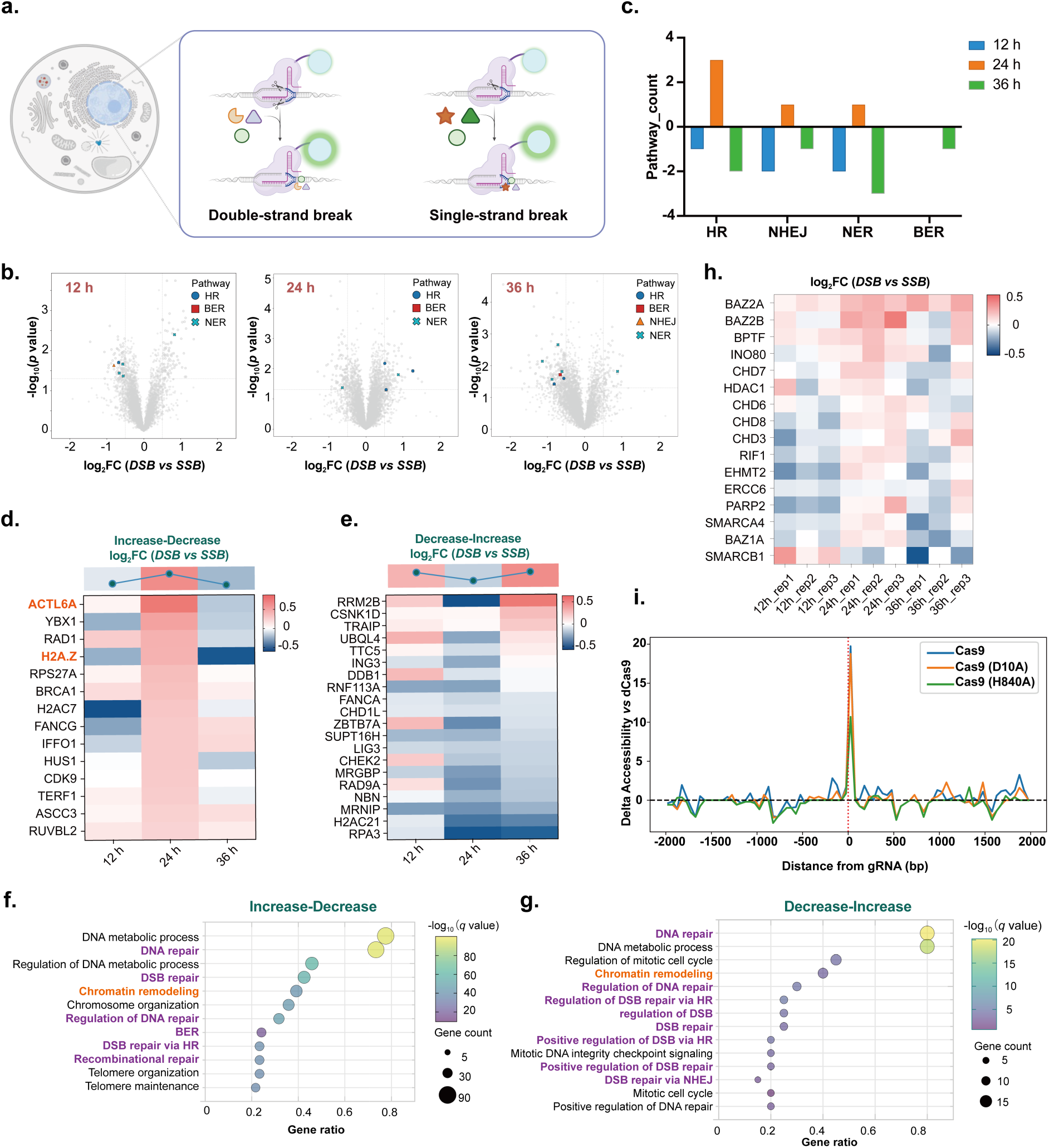
Cas9-TurboID for locus-specific DNA repair microenvironment labeling at the endogenous site. **a,** Scheme for DSB repair and SSB repair microenvironment labeling for proteomics analysis at the *EMX1* locus. **b,** Volcano analysis with repair pathways of DSB and SSB at 12 h, 24 h, and 36 h at the *EMX1* locus. Differentially significantly enriched proteins were defined using *p* < 0.05 and |log_2_FC| ≥ 0.5. **c,** Summary of pathway-associated proteins identified from the volcano plots, showing their distribution and counts across 12 h, 24 h, and 36 h conditions at *EMX1*. For each pathway, proteins with positive log₂FC values were classified as DSB-associated, and proteins with negative log₂FC values were classified as SSB-associated. **d-g,** Representative DNA repair-related proteins with different trends displaying distinct enrichment patterns between DSB and SSB conditions, together with corresponding Gene Ontology (GO) analysis at *EMX1*^36^. **d** and **f** represent Increase-Decrease protein analysis; **e** and **g** represent Decrease-Increase protein analysis. Data are presented as the average value of log_2_FC (DSB *vs* SSB) at 12 h, 24 h, and 36 h from three independent biological replicates (n = 3). **h,** Chromatin remodeling-related proteins enriched in DSB repair and SSB repair, shown as log_2_FC (DSB *vs* SSB) at 12 h, 24 h, and 36 h with three replicates. **i,** ATAC detection of DNA accessibility at the target site after transfection of Cas9 and derivatives, Cas9(D10A), Cas9(H840A), and dCas9 at 24 h.

Interestingly, in the dynamic repair-associated protein analysis, ACTL6A was enriched in DSB repair at 24 h with reduced enrichment at 36 h, which is the same as in the Luc assay. In contrast, the histone variant H2A.Z exhibited distinct temporal dynamics between exogenous and endogenous DNA repair (**Fig. 3d**, **3e)**. Given that RUVBL2 is a core component of the TIP60/SRCAP complexes involved in H2A.Z deposition at damaged chromatin^37,38^, the concurrent enrichment of RUVBL2 and H2A.Z in the endogenous case suggests activation of chromatin remodeling following Cas9-induced DNA break. Such remodeling may contribute to increased chromatin accessibility and facilitate the recruitment of downstream DNA repair machinery. Similarly, ACTL6A is one component of the SWI/SNF chromatin-remodeling complex^19^, which is involved in nucleosome repositioning and chromatin organization to facilitate the recruitment of DNA repair proteins. GO analysis further revealed that dynamically recruited repair-associated proteins at *EMX1* were highly enriched in chromatin-remodeling pathways, with peak enrichment occurring around 24 h after DSB induction **(Fig. 3f-h)**.

To directly assess whether local chromatin accessibility is altered during editing, we conducted ATAC-seq around the *EMX1* locus following Cas9 transfection. Active Cas9 and Cas9(D10A) induced pronounced local increases in DNA accessibility, whereas Cas9(H840A) showed a weaker effect and catalytically inactive dCas9 produced minimal changes **(Fig. 3i)**^39^. Furthermore, CUT&RUN-qPCR analysis demonstrated greater Cas9 occupancy at the endogenous *EMX1* locus compared to the exogenous target (**Extended Data Fig. 4b**), underscoring the critical requirement for dynamic chromatin remodeling during endogenous genome editing.

### ACTL6A and H2A.Z improve Cas9 efficiency especially in heterochromatin

Based on our finding that ACTL6A and H2A.Z are dynamically recruited during genome editing and associated with changes in local DNA accessibility, we hypothesized that these factors could be exploited to enhance Cas9 editing performance. We first evaluated the contribution of endogenous ACTL6A and H2A.Z to Cas9-mediated editing using shRNA-mediated knockdown (**Fig. 4a, 4b, 4d, 4e**). Depletion of either protein compromised Cas9 editing efficiency at the *EMX1* locus (**Fig. 4c**, **4f**). Interestingly, ACTL6A knockdown also led to a concurrent reduction in total H2A.Z protein levels (**Extended Data Fig. 5a, 5b**).

**Fig. 4.**
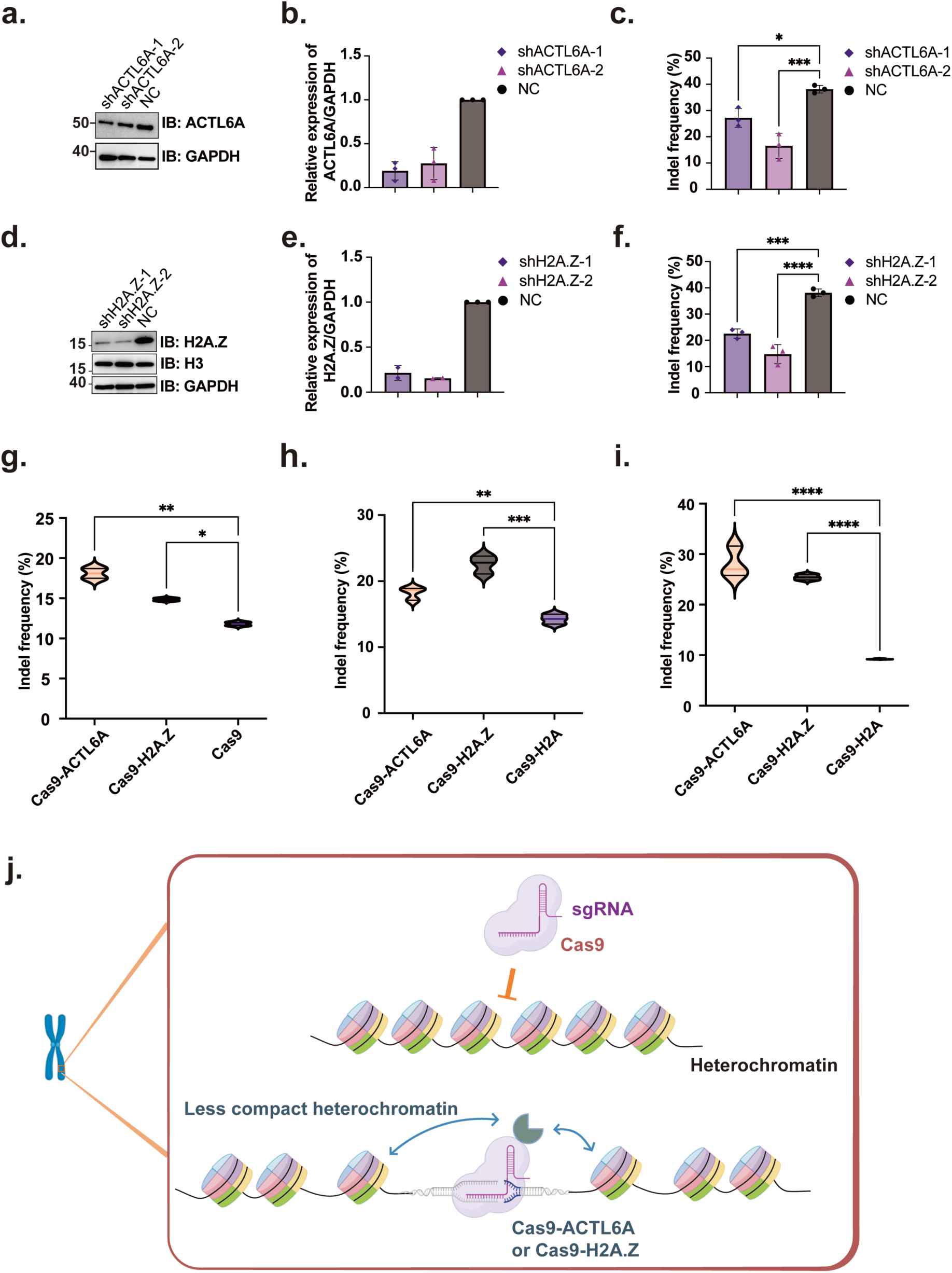
ACTL6A and H2A.Z affect the editing efficiency and help Cas9 editing in heterochromatin. **a,b,** Knockdown of ACTL6A by shRNA was validated by immunoblotting and quantified relative to the control. GAPDH served as a loading control. **c,** Depletion of ACTL6A reduced genome-editing efficiency at the *EMX1* locus, as determined by indel frequency detected by Sanger sequencing. Data are presented as mean ± s.d. from three independent biological replicates (n = 3). Statistical significance was determined using One-way ANOVA. *\*P* < 0.05, *\*\*P* < 0.01, *\*\*\*P* < 0.001. **d,e,** Knockdown of H2A.Z by shRNA was validated by immunoblotting and quantified relative to GAPDH. GAPDH and Histone 3 (H3) served as loading controls. **f,** Depletion of H2A.Z reduced genome-editing efficiency at the *EMX1* locus, as determined by indel frequency detected by Sanger sequencing. Data are presented as mean ± s.d. from three independent biological replicates (n = 3). Statistical significance was determined using One-way ANOVA. *\*P* < 0.05, *\*\*P* < 0.01, *\*\*\*P* < 0.001. **g-i,** Fusion of ACTL6A or H2A.Z to Cas9 enhanced editing efficiency. **g,** at the *EMX1* locus; **h,** at heterochromatic target locus on chromosomes 4 (Chr4); **i,** at heterochromatic target locus on chromosomes 17 (Chr17). Data are presented as mean ± s.d. from two independent biological replicates (n = 2) for the *EMX1* locus and three independent biological replicates (n = 3) for Chr4 and Chr17. Statistical significance was determined using One-way ANOVA. *\*P* < 0.05, *\*\*P* < 0.01, *\*\*\*P* < 0.001. **j,** Schematic illustration of heterochromatin editing mediated by Cas9-ACTL6A or Cas9-H2A.Z.

Inspired by previous studies demonstrating that Cas9-fused chromatin regulators can improve genome editing outcomes^40^, and given the roles of ACTL6A and H2A.Z in regulating chromatin organization and accessibility^41–43^, we engineered direct fusions of ACTL6A or H2A.Z to Cas9 and assessed their editing efficiency. Both Cas9-ACTL6A and Cas9-H2A.Z fusions enhanced editing efficiency at the canonical *EMX1* locus compared to wild-type Cas9 (**Fig. 4g**). Because heterochromatic regions are inherently refractory to genome editing due to tightly packed nucleosomes and limited Cas9 accessibility^44^, we next tested these fusions at heterochromatic genomic loci enriched in H3K9me3 (histone H3 lysine 9 trimethylation), identified using ENCODE databases^45^. Across two independent heterochromatic loci on chromosome 4 and chromosome 17 displaying varying H3K9me3 signal intensities, Cas9-ACTL6A and Cas9-H2A.Z fusions shows higher indel frequency compared to Cas9-H2A (**Fig. 4h**, **4i**). Previous studies have suggested that H2A.Z contributes to heterochromatin organization and interacts with HP1-mediated chromatin regulation^46^. The compact nature of heterochromatin limits Cas9 access to genomic DNA and constrains its target-binding dynamics. By locally remodeling the chromatin environment, Cas9-ACTL6A and Cas9-H2A.Z may increase nucleosome accessibility and facilitate Cas9 engagement at heterochromatic loci (**Fig. 4j**).

### Proteomic profiling of prime editing reveals enrichment of proteins recognizing complex DNA repair intermediates and identifies WRN as a functional regulator of prime editing

Beyond DSB and SSB repair, we further extended our spatiotemporal proximity-labeling platform to prime editing (PE), a versatile genome editing platform that enables precise sequence modifications without generating canonical double-strand breaks^4^. Compared to DSB and SSB, PE introduces substantially greater structural complexity by generating a series of transient repair intermediates, including RNA-DNA hybrids, heteroduplex DNA, and DNA flap structures, which require coordinated processing by multiple DNA repair pathways. Although recent studies have begun to identify factors influencing PE efficiency^47–49^, the local protein microenvironment associated with PE-mediated DNA processing remains largely unclear.

To systematically profile the protein microenvironments formed during prime editing, we engineered a prime editor-TurboID (PE-TurboID) fusion comprising nCas9(H840A) and a reverse transcriptase (RT). Functional validation confirmed that TurboID fusion did not compromise prime editing activity in either PE2 or PE3 (**Fig. 5a-5c**) while enabling robust biotinylation (**Extended Data Fig. 6a**). We selected the PE2 system, containing the prime editor and pegRNA, for downstream temporal analysis to minimize potential confounding effects introduced by the auxiliary nicking sgRNA in PE3. The editing efficiency gradually increased over time, reaching a plateau at 48 h **(Fig. 5d)**. Consistent with the editing kinetics observed above, 12 h, 24 h, and 36 h were selected as representative time points for profiling the dynamic assembly of PE-associated proteins (**Fig. 5e**). Quantitative pathway analysis revealed fewer canonical pathway-associated repair factors enriched in PE compared to DSB/SSB conditions (**Extended Data Fig. 6b, 6c**), reflecting that prime editing engages a distinct repair landscape associated with its unique DNA structures and processing intermediates.

**Fig. 5.**
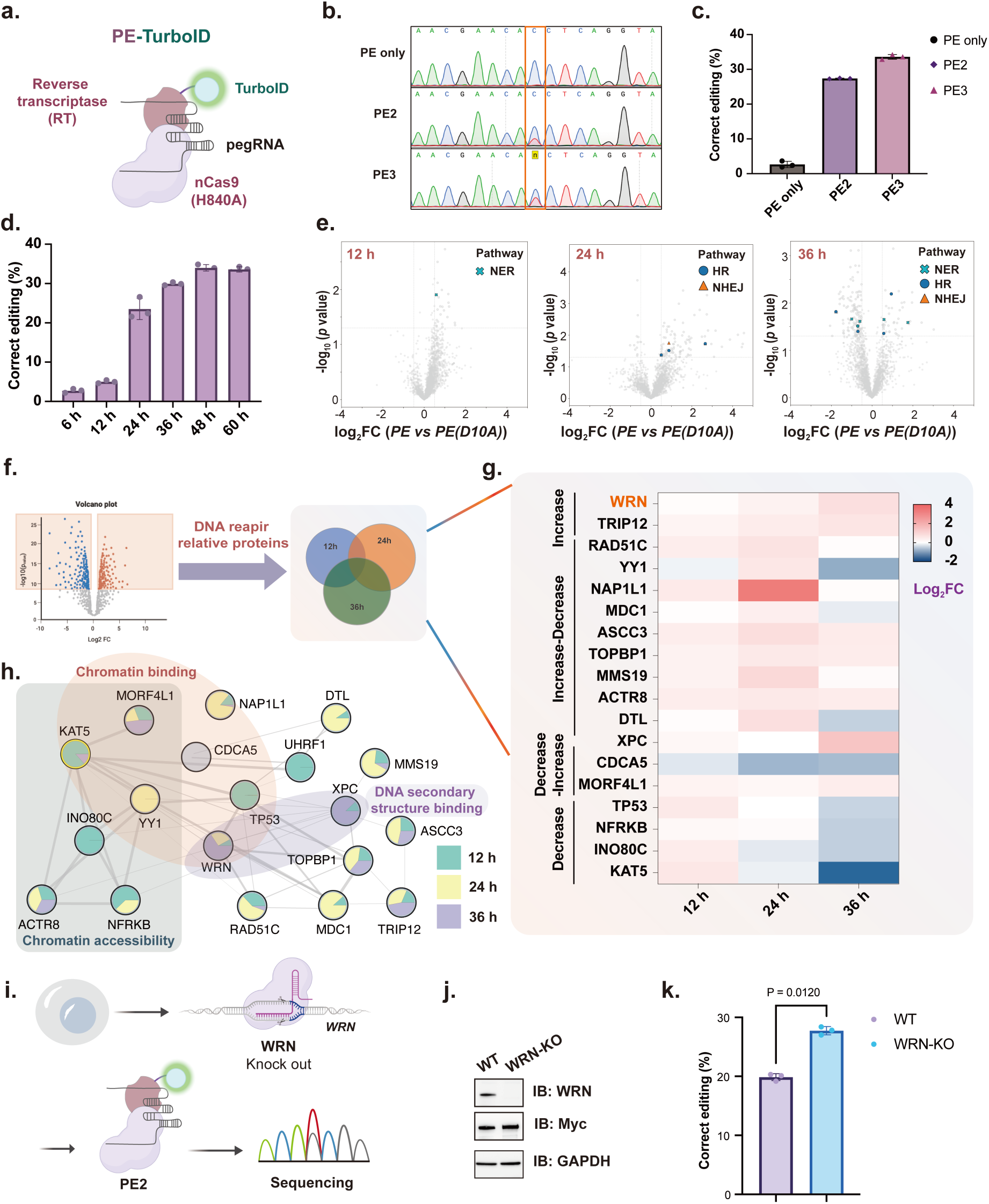
Proteomic mapping of the prime editing microenvironment. **a,** Schematic illustration of the construction of prime editor with TurboID as PE-TurboID. **b,c,** Editing activities of PE-TurboID detected by Sanger sequencing **(b)** and quantified as correct editing efficiency **(c).** Data are presented as mean ± s.d. from three independent biological replicates (n = 3). PE only, prime editor without pegRNA or sgRNA; PE2, prime editor with pegRNA; PE3, prime editor with pegRNA and sgRNA. **d,** Time-resolved analysis of correct editing efficiencies mediated by PE-TurboID, showing the dynamics of repair during prime editing. Data are presented as mean ± s.d. from three independent biological replicates (n = 3). **e,** Volcano plots showing proteins differentially enriched by PE-TurboID relative to PE(D10A)-TurboID at 12 h, 24 h and 36 h. Representative DNA repair pathways are indicated. Proteins were considered significantly differentially enriched when *P* < 0.05 and |log_2_FC (PE *vs* PE(D10A))| ≥ 0.5. **f,g,** Significantly enriched proteins, *P* < 0.05 and |log_2_FC| ≥ 0.5, at all time points were filtered as DNA repair proteins for further trend analysis, as Increase, Increase-Decrease, Decrease-Increase, and Decrease patterns according to the log_2_FC (PE *vs* PE(D10A)). **h,** Cluster map showing proteins enriched dynamically according to STRING analysis^61^ and shown with Cytoscape^62^. **i,** Schematic of the workflow for assessing prime editing efficiency in WRN-knockout cells. Wild-type (WT) and WRN-knockout (WRN-KO) HEK293T cells were transfected with PE-Myc, followed by assessment of prime editing efficiency at 36 h after transfection. **j,** Immunoblotting analysis confirming WRN knockout and expression of PE-Myc, with GAPDH as a loading control. **k,** Prime editing efficiency at the *RNF2* target site in WT and WRN-KO HEK293T cells. Data are presented as mean ± s.d. from three independent biological replicates (n = 3). Statistical significance was assessed using an unpaired two-sided Student’s *t*-test.

To define a core set of PE-associated machinery, we extracted DNA repair-related proteins consistently enriched across all three time points and categorized them into four dynamic temporal patterns: Increase, Increase-Decrease, Decrease-Increase, and Decrease (**Fig. 5f**). Temporal profiling of proteins recruited to prime editing sites revealed distinct recruitment dynamics during the editing process. Chromatin regulators, including KAT5 from the NuA4 complex and INO80C and ACTR8 from the INO80 complex, exhibited peak enrichment at early time points and subsequently declined, suggesting an early engagement of chromatin remodeling machinery (**Fig. 5g**, **5h**)^50–52^. The histone chaperone NAP1L1 peaked around 24 h, consistent with the continuous process of chromatin remodeling and histone binding^53,54^. In addition, DNA repair and replication stress-associated proteins, including RAD51C, TOPBP1, MMS19, and DTL, reached maximal enrichment at 24 h, indicating active engagement of repair-associated processes (**Fig. 5g**, **5h**)^55–58^.

Notably, structure-specific DNA-binding proteins displayed delayed or sustained enrichment kinetics, marked by the progressive accumulation of the RecQ helicase WRN and the late-stage recruitment of XPC at 36 h. Cluster analysis identified WRN and XPC within a DNA secondary structure-binding module^59–62^, consistent with the presence of structurally complex DNA intermediates associated with prime editing (**Fig. 5g**, **5h**)^63^. Collectively, these results show that proximity labeling captures the sequential recruitment of chromatin remodelers, replication stress responders, and DNA structure-associated factors during prime editing.

Next, we investigated whether the identified factors could functionally influence prime editing efficiency. We selected WRN, a DNA structure-associated factor with helicase and exonuclease activities, for functional validation. After generating a WRN-knockout (WRN-KO) cell line, PE2 editing efficiency was assessed (**Fig. 5i**). Immunoblotting confirmed WRN knockout and PE2 editor expression (**Fig. 5j**). At the *RNF2* target site, PE2 editing efficiency improved from 20% to 27% in WRN-KO cells compared to wild-type cells (**Fig. 5k**), identifying WRN as a potential regulator of prime editing.

## Discussion

In this study, we established a temporally resolved proximity-labeling platform by directly fusing genome editors with the proximity-labeling enzyme TurboID, enabling systematic mapping of the local protein microenvironments assembled during genome editing. By comparing exogenous target and endogenous genomic locus, we revealed that DNA repair-associated factors display both conserved and context-dependent recruitment patterns, highlighting the critical role of chromatin architecture in shaping repair assembly. Specifically, Cas9 exhibited increased genomic occupancy at the endogenous *EMX1*, accompanied by the robust 24 h recruitment of chromatin remodelers and a corresponding increase in local DNA accessibility measured by ATAC-seq. These findings reveal that endogenous repair occurs within a dynamic chromatin environment, where coordinated chromatin remodeling facilitates access to nucleosome-constrained genomic regions.

Through dynamic comparative profiling between DSB and SSB repair pathways, we identified two proteins, ACTL6A and H2A.Z, that exhibited distinct kinetic recruitment patterns. Functional depletion of either factor compromised Cas9 efficiency, whereas direct fusion of ACTL6A or H2A.Z to Cas9 substantially enhanced editing outcomes at *EMX1* and in heterochromatin, which is typically refractory to genome editing due to limited DNA accessibility. Consistent with their known roles in chromatin remodeling, both ACTL6A and H2A.Z may enhance Cas9 editing efficiency by facilitating access to genomic DNA.

Extending this proximity-labeling framework to prime editing reveals that prime editing involves a series of DNA intermediates that require coordinated processing by chromatin regulatory and DNA repair pathways. Consistent with this concept, we observed temporally distinct recruitment patterns among chromatin regulators and DNA secondary structure-binding proteins. Early enrichment of KAT5 and INO80 complex components was followed by accumulation of repair-associated proteins such as RAD51C and TOPBP1, whereas WRN and XPC exhibited sustained or delayed enrichment profiles. These observations suggest that different cellular pathways may be engaged at different stages of the editing process. Notably, NAP1L1, WRN, and XPC displayed the most prominent temporal dynamics. Although the functional significance of their recruitment remains to be determined, their enrichment patterns highlight them as candidate regulators of prime editing-associated cellular responses. Following candidate selection, we functionally validated WRN as a regulator of prime editing and found that WRN knockout enhanced the editing efficiency of PE2. Previous studies have implicated WRN activity in the response to mismatch repair (MMR) deficiency and microsatellite instability (MSI)^64^, suggesting a role for WRN in the processing of aberrant DNA structures in these cellular contexts. Given that prime editing involves the formation and resolution of distinct DNA intermediates, our finding raises the possibility that WRN-associated DNA processing may influence prime editing outcomes. Whether this effect is mediated by the helicase activity of WRN or other functions of the protein remains to be determined. Further investigation of how WRN and related DNA-processing factors act on prime-editing intermediates may provide additional insight into the cellular determinants of prime editing efficiency.

Overall, this study establishes a temporally resolved proximity-labeling framework that is broadly applicable across diverse genome editing modalities. Beyond identifying ACTL6A and H2A.Z as engineering modules for enhancing genome editing, this framework provides a systematic resource for characterizing the local molecular environment of programmable editors in different cellular and chromatin contexts. We further demonstrate the utility of this approach for identifying functional regulators of prime editing, exemplified by the enhanced prime editing efficiency observed following WRN knockout. Such knowledge may facilitate the rational design of genome editors with improved efficiency, specificity, and context-dependent performance, while providing a versatile platform for mechanistic studies and future editor engineering. More broadly, the molecular environments captured by this platform provide a foundation for future investigations of locus-specific DNA damage responses, DNA repair processes, and other genome-associated events within their native chromatin contexts. This method can be applied to a broad scope of genome editors.

## Methods

### General molecular biology and cloning

DNA amplification was conducted by PCR using Phanta Flash Master Mix (Vazyme, P510). Primers were synthesized by Genewiz (Azenta Life Sciences, Suzhou, China). For plasmid construction, corresponding sequences were PCR-amplified and cloned into the indicated expression vectors using Gibson assembly. The prime editing plasmid of PE2 was constructed based on Addgene 174828 by deleting the hMLH1dn sequence. Plasmids expressing pegRNAs and sgRNAs were constructed by Gibson assembly or Golden Gate assembly based on Addgene 132777. pegRNA and sgRNA sequences are shown in **Supplementary Table 2**. TurboID plasmid (Addgene 107169) was amplified by PCR and cloned into Cas9 and PE. ACTL6A and H2A.Z were synthesized based on the NM_004301.5 and NM_002106.4, respectively.

### Cell culture

HEK293T cells (catalogue number 1101HUM-PUMC000091) were purchased from the Cell Resource Center, Peking Union Medical College, China. HEK293T cells were maintained in DMEM (Dulbecco’s modified Eagle’s medium, Gibco) containing 10% fetal bovine serum (Gibco) and 1% penicillin/streptomycin (Gibco), and cultured at 37 °C under 5% CO_2_.

### Transfection for plasmid DNA delivery

HEK293T cells were seeded in 24-well plates for 16-24 h until approximately 50-60% confluency. For the luciferase reporter assay, 500 ng of Cas9-Luc plasmid and 500 ng of fLuc-TAG plasmid were co-transfected with Lipofectamine 3000 (Invitrogen) according to the manufacturer’s protocol. To perform genome editing at the endogenous *EMX1* locus, 1000 ng of Cas9-EMX1 plasmid was transfected under the same conditions described above. For PE, 1500 ng PE plasmid, 500 ng pegRNA, and 166 ng sgRNA plasmid (for PE3) were co-transfected under the same conditions described above.

### Genomic DNA extraction and amplification for sequencing

Cells were collected and washed with PBS three times. The genomic DNA mixture was incubated in fresh lysis buffer (10 mM Tris-HCl, pH 7.5, 0.05% SDS, and 400 μg/ml proteinase K) at 37 °C for 1.5 h, followed by an 80 °C deactivation step for 30 min. Targeted genomic regions were amplified using PCR^4^. The thermal cycling conditions were set at 95 °C for 3 min; 30 cycles of 95 °C for 10 s, 56-60 °C for 10 s, and 72 °C for 1 min; followed by a final extension of 72 °C for 5 min. Primers used in this paper are listed in **Supplementary Table 3**. Primers were screened by BLAST to ensure target specificity, and the corresponding genomic targets are listed in **Supplementary Table 4**.

### Sequencing analysis

Sanger sequencing data for editing were analyzed by Tider (https://tide.nki.nl).

### Biotin labeling in HEK293T cells

HEK293T cells were transfected with Cas9-TurboID or Cas9(D10A)-TurboID in *EMX1* case or an additional Luc plasmid in the *Luc* case for the indicated time. In the PE case, cells were transfected with PE-TurboID or PE(D10A)-TurboID with an additional pegRNA plasmid for the indicated time. Then the medium was replaced with 500 μM biotin in prewarmed serum-containing cell culture medium for 20 min^18^. Labeling was stopped after the desired time period by transferring the cells to ice and washing three times with ice-cold PBS.

### Streptavidin enrichment and on-bead digestion for LC–MS/MS analysis

Labeled cells were collected at 500 g for 3 min. The supernatant was removed, and the pellet was lysed in RIPA lysis buffer (Thermo Fisher Scientific, 89900) by gentle pipetting and sonicated on ice. Lysates were clarified by centrifugation at 12,000 g for 10 min at 4 °C. Streptavidin magnetic beads (Thermo Fisher Scientific, 20359) were pre-washed three times with PBS and incubated with the lysates for 3 h at room temperature. Beads were subsequently washed once with 0.2% SDS/PBS, three times with PBS, and three times with distilled water.

For on-bead digestion, proteins were reduced in 6 M urea in PBS containing 10 mM dithiothreitol at 37 °C for 30 min and alkylated with 20 mM iodoacetamide at 35 °C for 30 min in the dark. Beads were washed once with 2 M urea/PBS and resuspended in 2 M urea/PBS containing sequencing-grade trypsin. Digestion was performed overnight at 37 °C with shaking. Peptides were collected from the supernatant and additional water washes, then concentrated using a Speed-Vac concentrator.

Peptides were desalted using C18 spin columns. Columns were activated with 50% acetonitrile, equilibrated with 0.1% formic acid (FA), and loaded with peptide samples dissolved in 0.1% FA. After washing with 0.1% FA followed by 0.1% FA in 5% acetonitrile, peptides were eluted with 0.1% FA in 90% acetonitrile. Eluates were dried in a Speed-Vac concentrator and subjected to LC-MS/MS analysis in data-independent acquisition (DIA) mode.

### Analysis of mass spectrometry data

Raw DIA-MS data were processed using DIA-NN (v1.8.1) in library-free mode against the UniProt human proteome database. Peptide and protein identifications were filtered at a 1% false discovery rate (FDR). Protein quantification was performed using the MaxLFQ algorithm implemented in DIA-NN. The resulting protein intensity matrix was used for downstream statistical analyses.

### Confocal fluorescence imaging of cultured cells

HEK293T cells were plated, transfected, and labeled with biotin as described above, and subsequently fixed with 4% (v/v) paraformaldehyde (Beyotime) in PBS at room temperature for 15 min. Cells were then washed three times with PBS and permeabilized with 0.25% Triton -X-100 (Beyotime) in PBS at room temperature for 15 min. Cells were then washed three times with PBST and incubated with 5% skim milk in PBST at room temperature for 1 h, and then washed with PBST three times, followed by primary antibody (**Supplementary Table 1**) in PBST at room temperature for 1 h or 4 °C overnight. After washing three times with PBST, cells were then incubated with Hoechst 33342/ fluorochrome-conjugated secondary antibody (**Supplementary Table 1**) in PBST for 1 h at room temperature. Cells were then washed three times with PBST and imaged by confocal fluorescence microscopy.

### Construction of the knockdown cell line with shRNA

For preparation of lentiviruses, HEK293T cells in 12-well plates were transfected at ∼70% confluency with the pLKO.1-shRNA (sequence was shown in **Supplementary Table 5)**, the lentiviral packaging plasmids pMD2.G and psPAX2, and Polyethylenimine (Sigma-Aldrich) for 8 hours. The medium was exchanged with fresh medium. About 48 h after transfection, the cell culture medium was harvested and the lentiviruses mixed with 8.5 µg/mL polybrene (Merck) and incubated with target cells. HEK293T cells seeded in a 6-well plate were then infected at 50% confluency, followed by selection with 2 µg/mL puromycin (Harveybio) in culture medium for a week before further analysis.

### Construction of the knockout cell line

HEK293T cells were transfected with the PX458 plasmid (Addgene 48138) encoding Cas9-EGFP with a guide RNA targeting the *WRN* gene (gRNA sequence: 5′-GCATCGTAACTATACACAA-3′). At 48 h after transfection, GFP-positive single cells were isolated by fluorescence-activated cell sorting (FACS) and expanded to establish individual clonal cell lines. Clones were screened by Sanger sequencing and confirmed for loss of WRN protein expression by immunoblotting. A validated knockout clone was used for subsequent experiments.

### Immunoblotting analysis

For immunoblotting analysis, samples were separated through SDS-PAGE and transferred to 0.45 or 0.2 μm PVDF membrane (Millipore, IPVH00010 or ISEQ00010). Membranes were blocked with 5% skim milk dissolved in TBST, then incubated with primary antibodies and secondary antibodies at appropriate concentrations (dissolved in antibody dilution buffer, Epizyme, PS119L) sequentially. All the immunoblot images were captured by Chemdoc Bio-Rad.

### CUT&RUN-qPCR

CUT&RUN–qPCR was performed using the CUT&RUN Assay Kit for qPCR (Vazyme, HD103) according to the manufacturer’s instructions. Briefly, approximately 1 × 10^5^ cells were harvested after being transfected with Cas9-Flag for the indicated time and immobilized on ConA-coated magnetic beads. Cells were permeabilized and incubated overnight at 4 °C with anti-FLAG antibody (Cell Signaling Technology, 14793S) diluted 1:100 in Antibody Buffer. Following antibody binding, samples were washed with digitonin-containing wash buffer and incubated with pA/G-MNase at 4 °C for 1 h. Excess pA/G-MNase was removed by washing, and chromatin digestion was initiated by the addition of CaCl₂. Digestion was performed on ice for 90 min and terminated with Stop Buffer containing spike-in DNA. Cleaved chromatin fragments were released by incubation at 37 °C for 30 min. DNA was subsequently purified using the DNA extraction beads provided in the kit and eluted in nuclease-free water.

Quantitative PCR was performed using SupRealQ Ultra Hunter SYBR qPCR Master Mix (U+) (Vazyme, Q713) according to the manufacturer’s instructions. Each reaction contained purified CUT&RUN DNA and locus-specific primers in **Supplementary Table 3**. Amplification was carried out under the following cycling conditions: 95 °C for 30 s, followed by 40 cycles of 95 °C for 10 s, 56 °C for 30 s, and 72 °C for 30 s. Melting curve analysis was performed at 95 °C for 15 s, 60 °C for 1 min, and 95 °C for 15 s. Relative enrichment was determined by qPCR and normalized to the corresponding spike-in DNA.

### ATAC-seq library preparation and sequencing

ATAC-seq libraries were generated using the Hyperactive ATAC-Seq Library Prep Kit for Illumina (Vazyme, TD711) according to the manufacturer’s instructions. Briefly, cells transfected with the corresponding plasmid (Cas9-Flag, Cas9(D10A)-Flag, Cas9(H840A)-Flag, and dCas9-Flag) were harvested and washed once with cold PBS. Nuclei were isolated in lysis buffer and incubated on ice for 10 min. The transposition reaction was performed using Tn5 transposase at 37°C for 30 min. Following transposition, reactions were terminated using stop buffer and DNA fragments were purified with ATAC DNA Extract Beads supplied in the kit.

Purified DNA was subjected to PCR amplification using indexed primers (Vazyme, TD202) and the amplification mix provided in the kit. Libraries were amplified under the following conditions: 72°C for 3 min, 95°C for 3 min, followed by 12 cycles of 98°C for 10 s and 60°C for 5 s, with a final extension at 72°C for 1 min. Amplified libraries were purified and size-selected using ATAC DNA Clean Beads (Vazyme, TD711). Library quality and fragment size distribution were assessed using an Agilent 2100 Bioanalyzer. Sequencing libraries were pooled and subjected to paired-end sequencing (2 × 150 bp) on an Illumina NovaSeq platform.

## Data availability

All data supporting the findings of this study are available within the Article, the Supplementary Information, and the Source data. The mass spectrometry proteomics data have been deposited to the ProteomeXchange Consortium (https://proteomecentral.proteomexchange.org) via the iProX partner repository^65,66^ with the dataset identifier PXD084318.

## Acknowledgements

We thank the National Center for Protein Sciences (Beijing) - Protein and Gene Platform at Peking University for the shRNA experiment. We thank the flow cytometry Core at National Center for Protein Sciences at Peking University, particularly Ms. Jia Luo, for technical help.

This work was supported by the Ministry of Science and Technology of China (2024YFA0917503, 2023YFA1506501, 2021YFA1302603 and 2022YFA1304700), the National Natural Science Foundation of China (22137001, 91753000 to P.R.C., 22222701, 22321005, 92253301 to X.F.), and the New Cornerstone Science Foundation through the New Cornerstone Investigator Program and the XPLORER PRIZE (P.R.C.).

## Author contributions

P.R.C. and X.F. conceived and supervised the project. S.X. performed the experiments and conducted the proteomic analysis. L.K. helped with proteomic experiments and knockdown experiments. W.Z. and X.L. helped with proteomic experiments. S.X., P.R.C., and X.F. wrote the manuscript with input from all authors.

---

**Extended Data Fig. 1.**
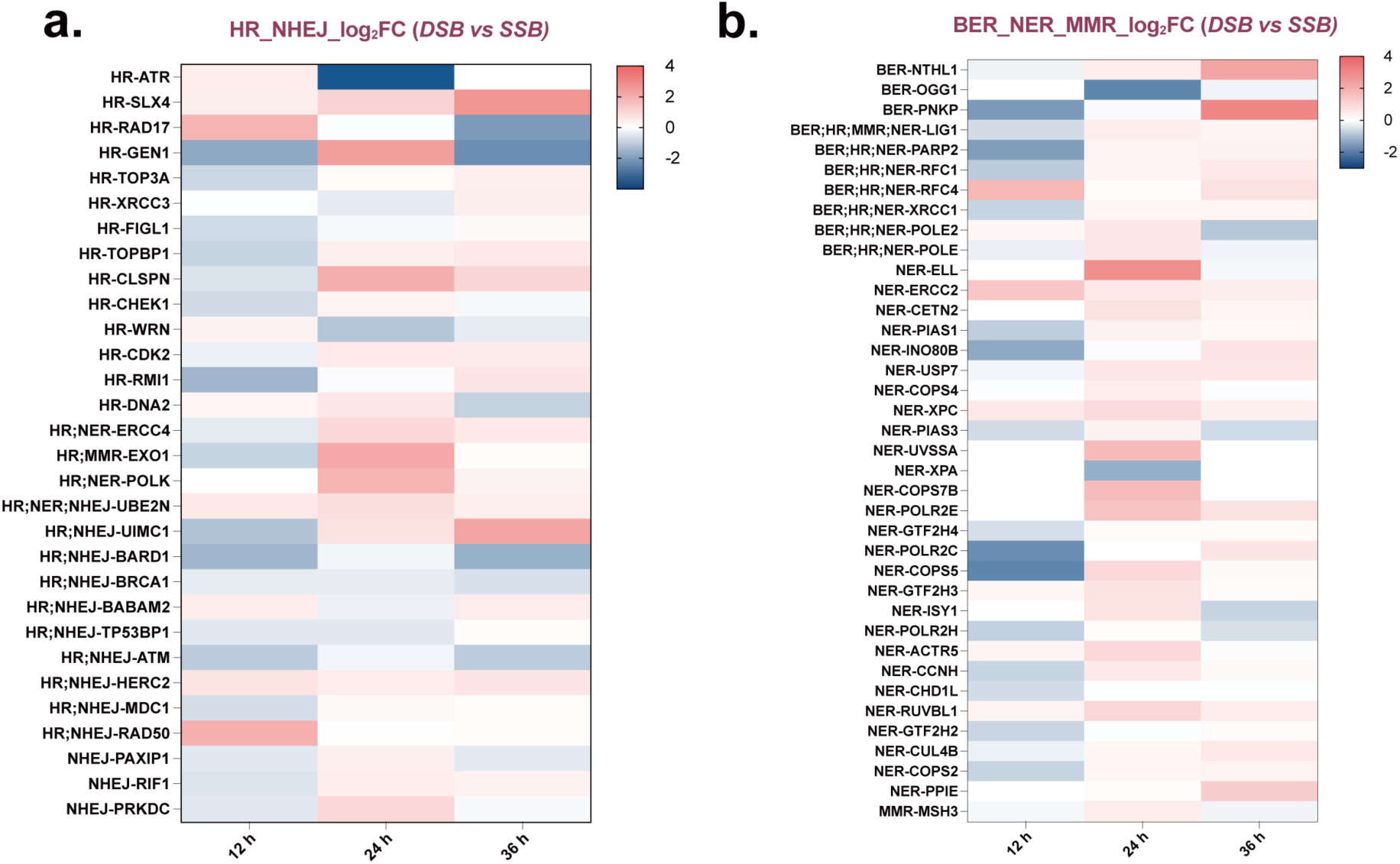
Proteomic analysis of proteins for Luc assay, classified by pathway. **a,** HR and NHEJ; **b,** BER, NER and MMR. Data are presented as the average value of log_2_ fold change of DSB *vs* SSB at 12 h, 24 h, and 36 h from three independent biological replicates (n = 3).

**Extended Data Fig. 2.**
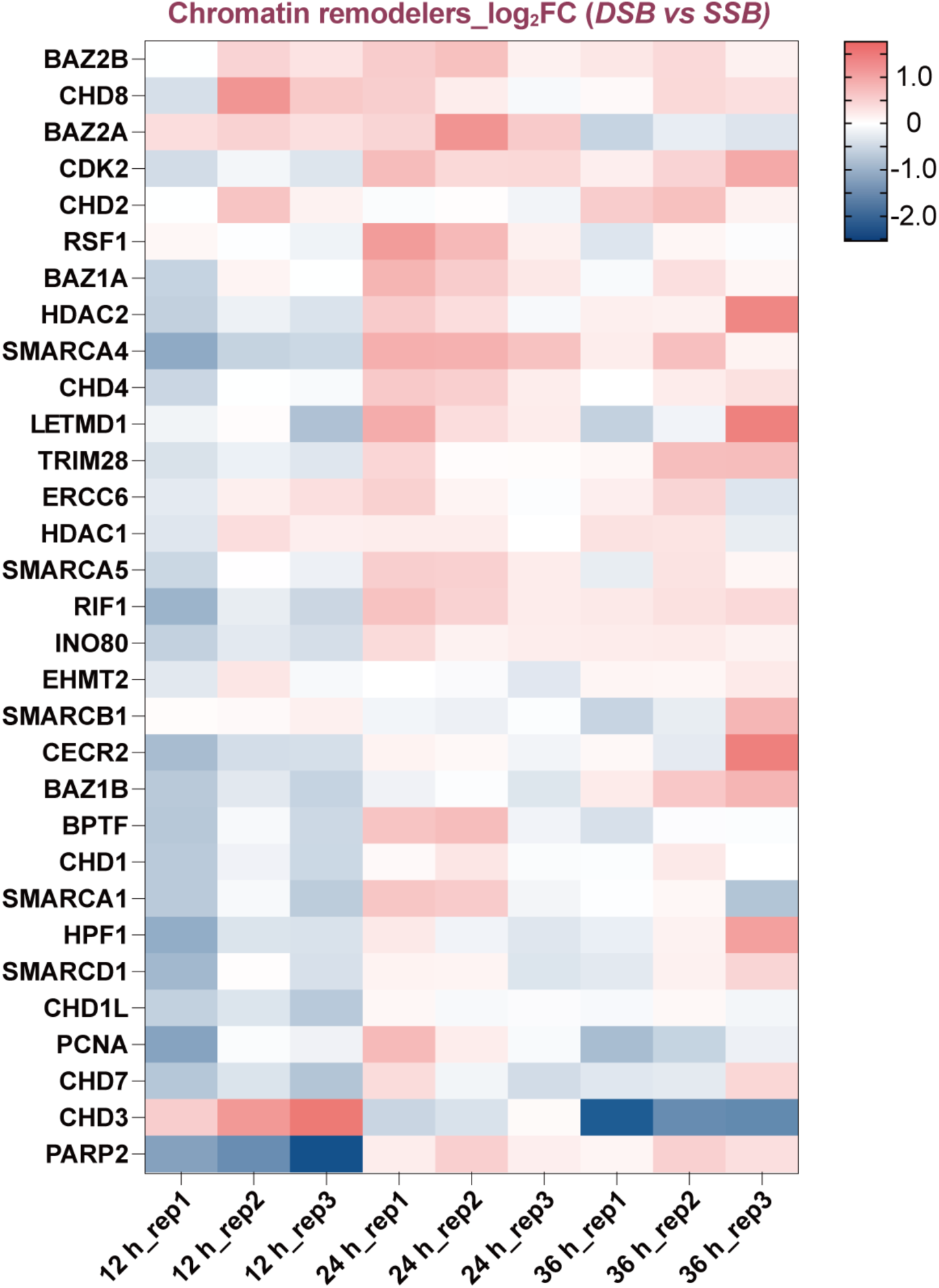
Dynamic changes in chromatin remodeling-related proteins following DSB and SSB induction in Luc assay. Data are presented as log_2_ fold change of DSB *vs* SSB at 12 h, 24 h, and 36 h, with three biological replicates per condition.

**Extended Data Fig. 3.**
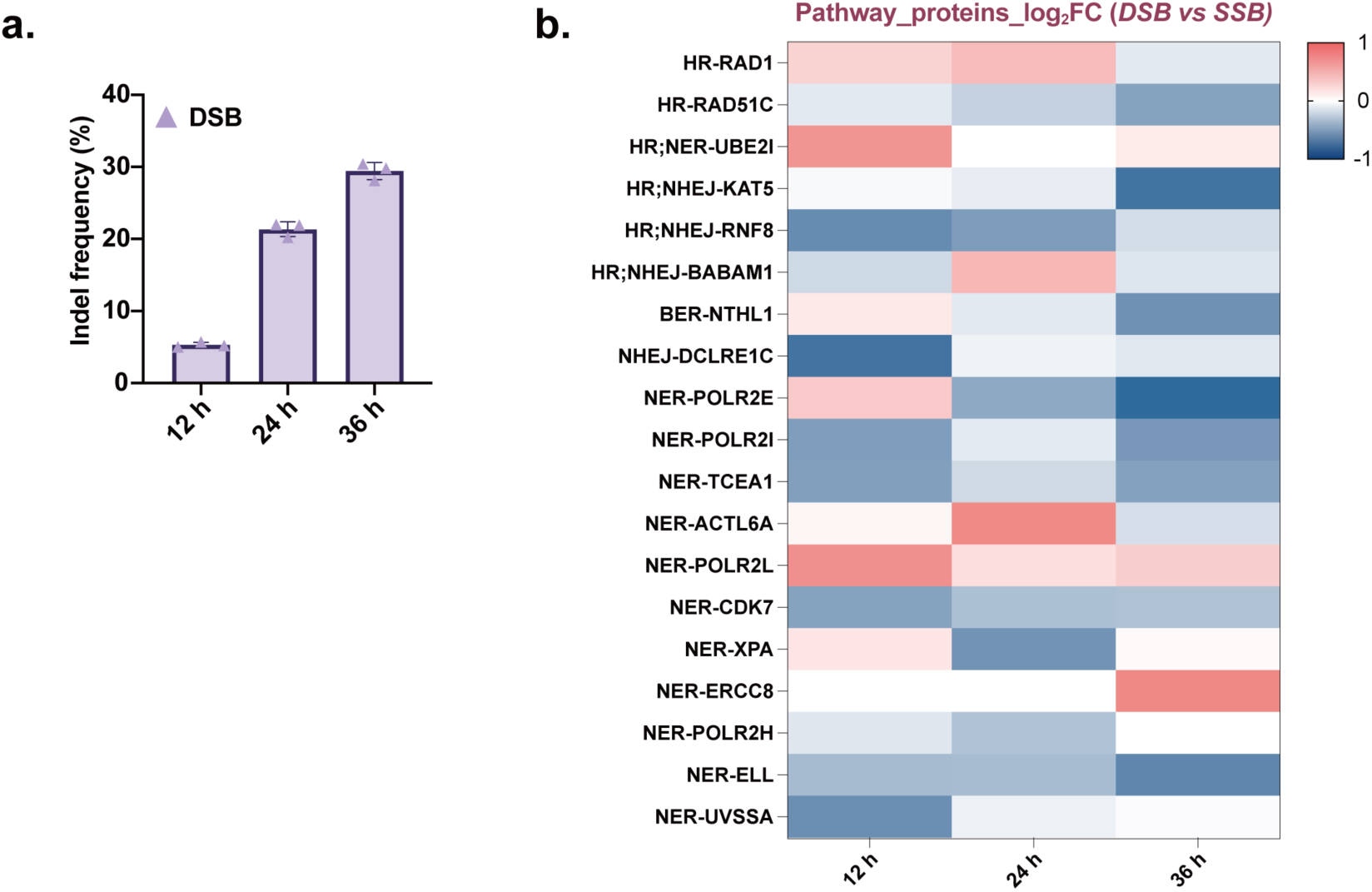
**a,** Cas9 editing at the *EMX1* site at 12 h, 24 h, and 36 h. Data were obtained by Sanger sequencing. Data are presented as mean ± s.d. from three independent biological replicates (n = 3). **b,** Proteomic analysis of proteins for *EMX1* assay, classified by pathway. Data are presented as the average value of log_2_ fold change of DSB *vs* SSB at 12 h, 24 h, and 36 h from three independent biological replicates (n = 3).

**Extended Data Fig. 4.**
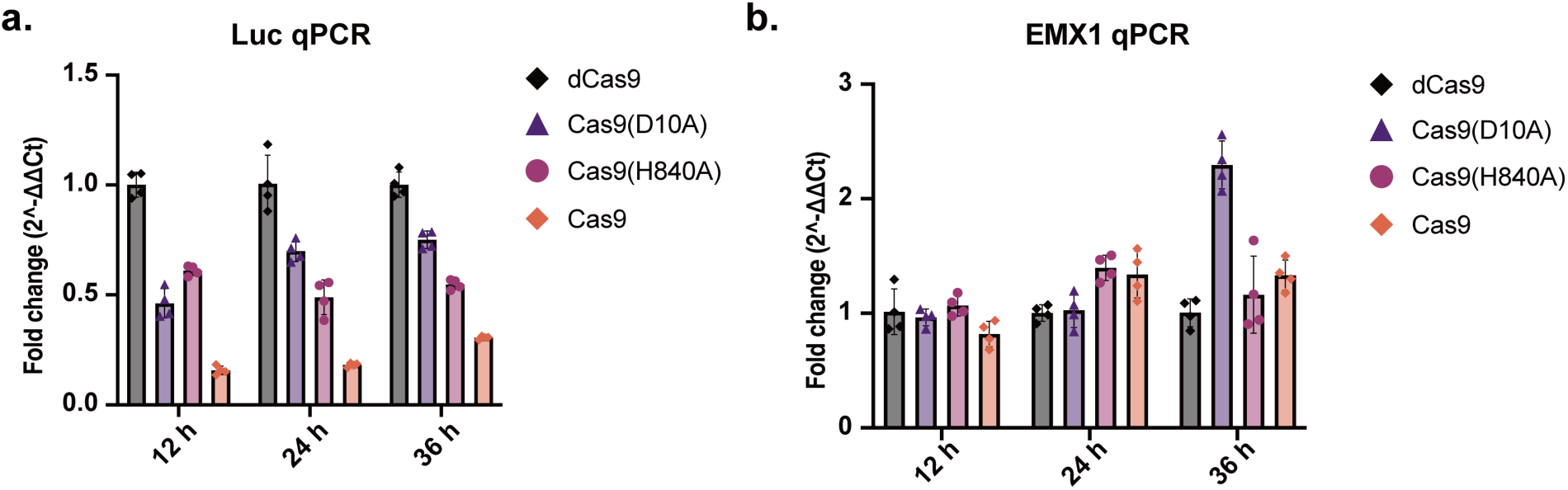
CUT&RUN-qPCR for quantification of enriched gRNA target sequence at 12 h, 24 h, and 36 h with wild-type Cas9, Cas9(D10A), Cas9(H840A), and dCas9. **a,** Luc assay. **b,** *EMX1* locus. Data are presented as mean ± s.d. from four independent biological replicates (n = 4).

**Extended Data Fig. 5.**
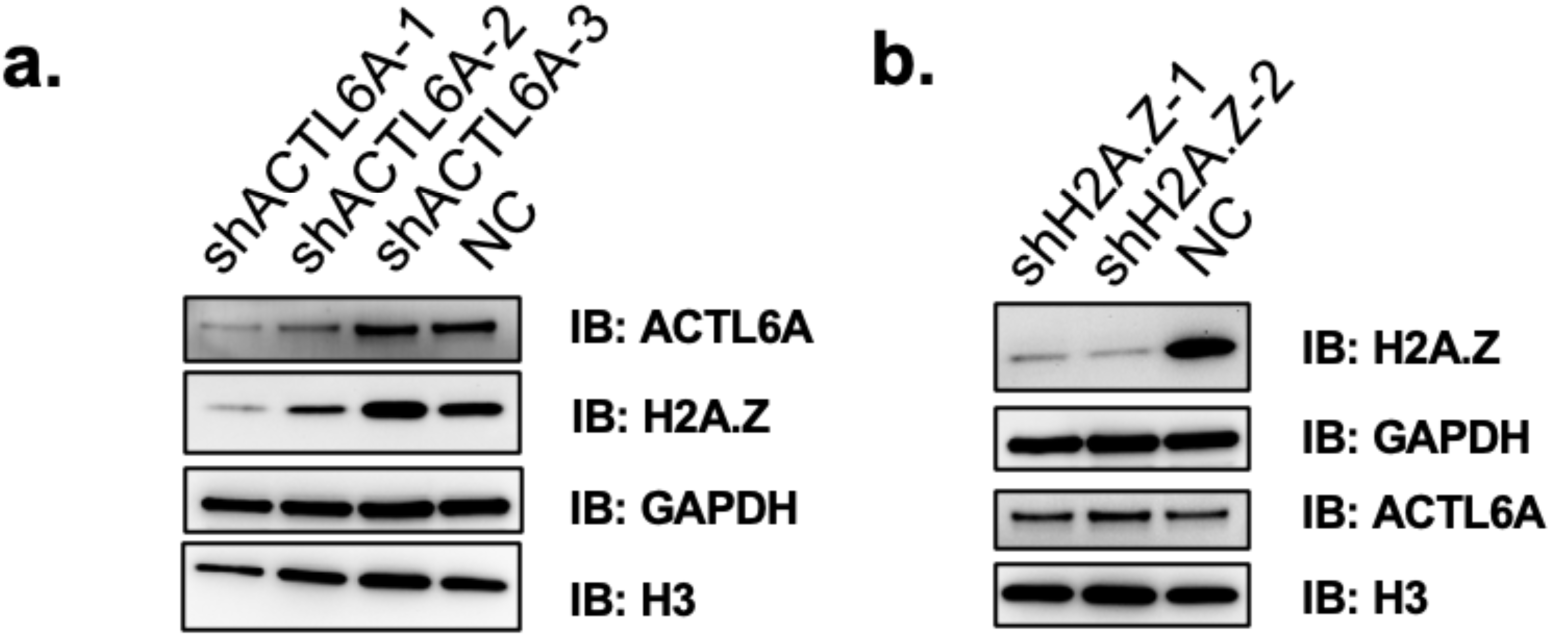
Knockdown of ACTL6A reduced the expression level of H2A.Z, **a**; while knockdown of H2A.Z did not affect the ACTL6A level, **b**. ACTL6A and H2A.Z protein levels in the knockdown cell lines were assessed by immunoblotting. GAPDH and Histone 3 (H3) were used as loading controls for total protein and nuclear protein, respectively.

**Extended Data Fig. 6.**
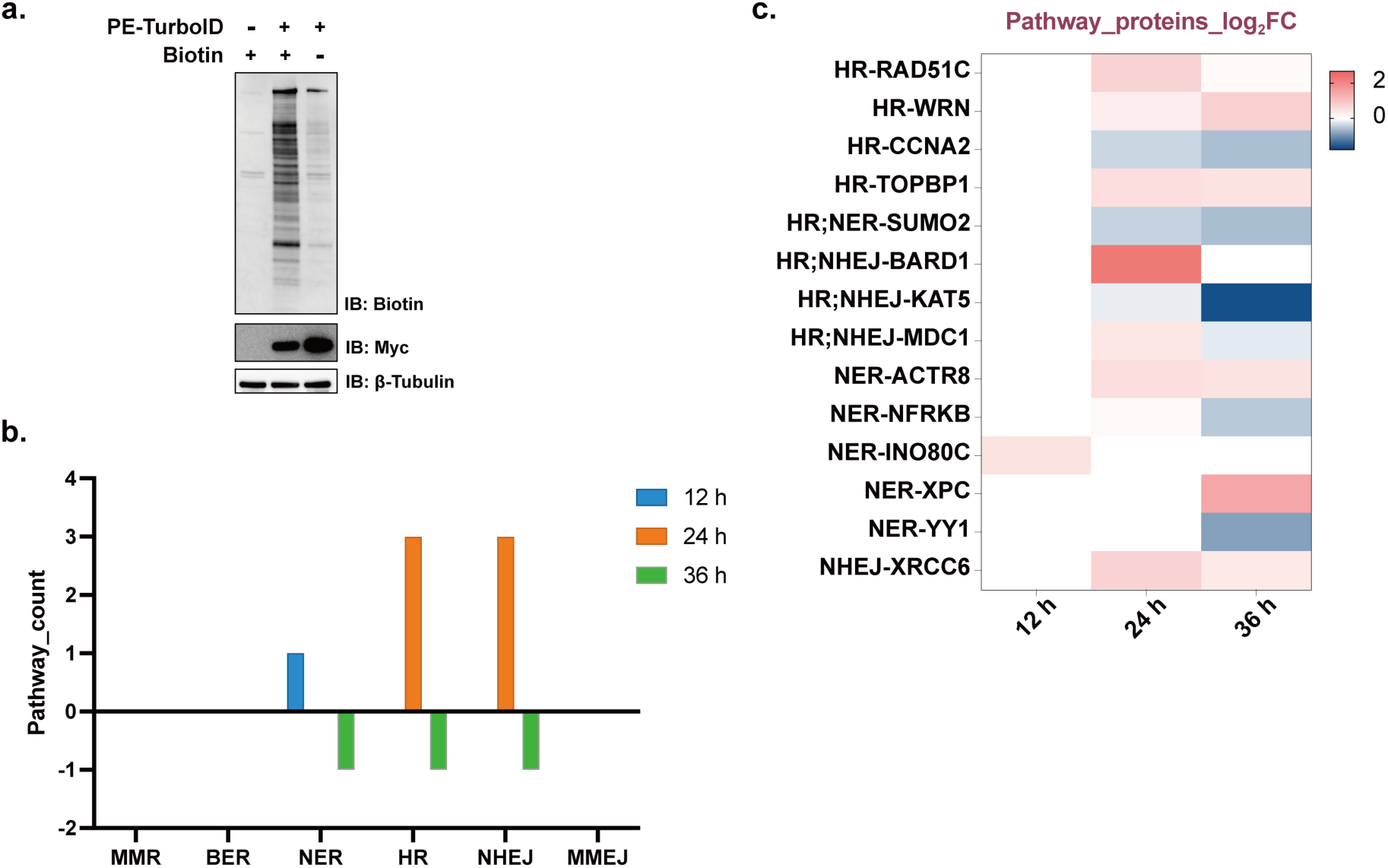
**a,** Immunoblotting analysis of biotinylated proteins demonstrating efficient TurboID-mediated proximity labeling in PE-TurboID-expressing cells. PE-TurboID-Myc was used for labeling, and β-Tubulin was used as a loading control. **b,** Count of enriched pathway proteins in PE. For each pathway, proteins with positive log₂FC (PE *vs* PE(D10A)) values are classified as recruitment in PE. **c,** Significantly enriched pathway proteins are shown with the corresponding pathway. Data are presented as the average value of log_2_ fold change of PE *vs* PE(D10A) at 12 h, 24 h, and 36 h from three independent biological replicates (n = 3).

**Supplementary Table 1.** Reagents.

| Reagent | Target species | Supplier | Cat. No. |
| --- | --- | --- | --- |
| ACTL6A polyclonal antibody | Rabbit | Proteintech | 10341-1-AP |
| Histone H2A.Z antibody | Rabbit | CST | 2718S |
| WRN antibody | Mouse | CST | 4666T |
| DYKDDDDK Tag antibody | Rabbit | CST | 14793S |
| DYKDDDDK Tag antibody | Mouse | CST | 8146S |
| Histone H3 antibody | Rabbit | CST | 4499S |
| GAPDH antibody | Mouse | CST | 97166S |
| Proteinase K | N/A | Harveybio | SR4920 |
| Puromycin | N/A | Harveybio | CT0049001 |
| Polybrene | N/A | Merck | TR-1003-G |
| Polyethylenimine branched (PEI) | N/A | Sigma-Aldrich | 408727 |
| RIPA buffer | N/A | Thermo Fisher Scientific | 89900 |
| 2 × Phanta Flash Master Mix | N/A | Vazyme | P510-03 |
| SupRealQ Ultra Hunter SYBR qPCR Master Mix(U+) | N/A | Vazyme | Q713-02 |
| Hoechst 33342 | N/A | Beyotime | C1027 |
| 4% (v/v) paraformaldehyde | N/A | Beyotime | P0099 |
| Triton-X100 | N/A | Beyotime | ST797 |
| Pierce High Capacity Streptavidin Agarose | N/A | Thermo Fisher Scientific | 20359 |

**Supplementary Table 2.**
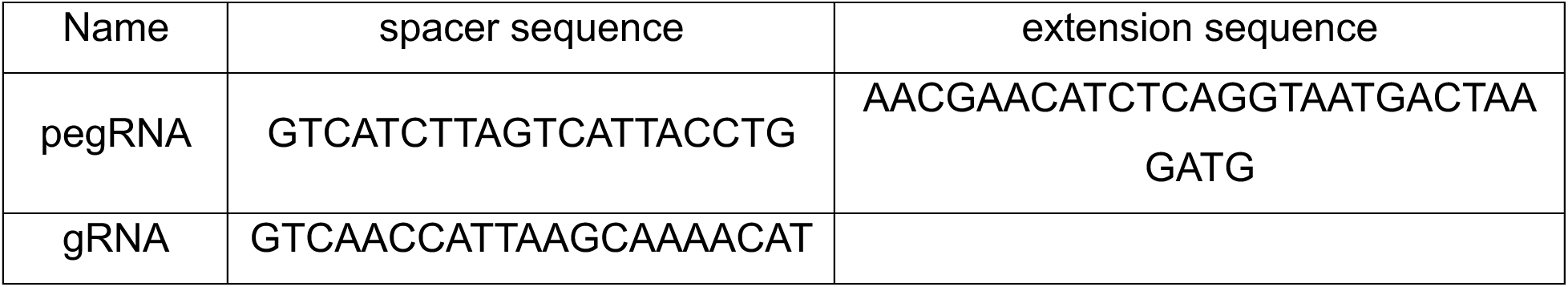
pegRNA used in PE2 and additional gRNA in PE3 at *RNF2*.

**Supplementary Table 3.**
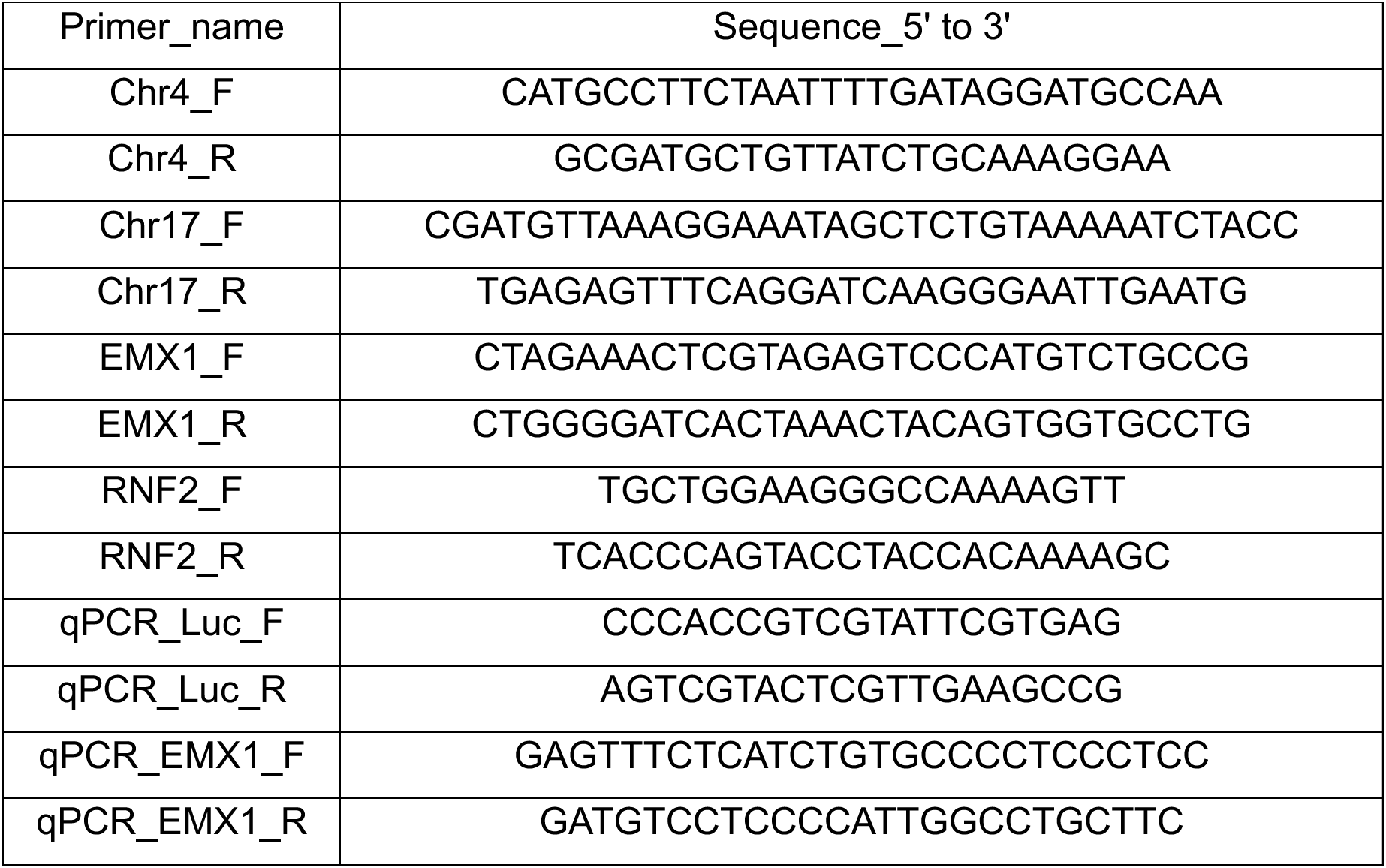
Primers used in this study. All sequences are shown in 5’ to 3’ orientation, and primer sequences were subjected to NCBI BLAST analysis to confirm target specificity.

**Supplementary Table 4.**
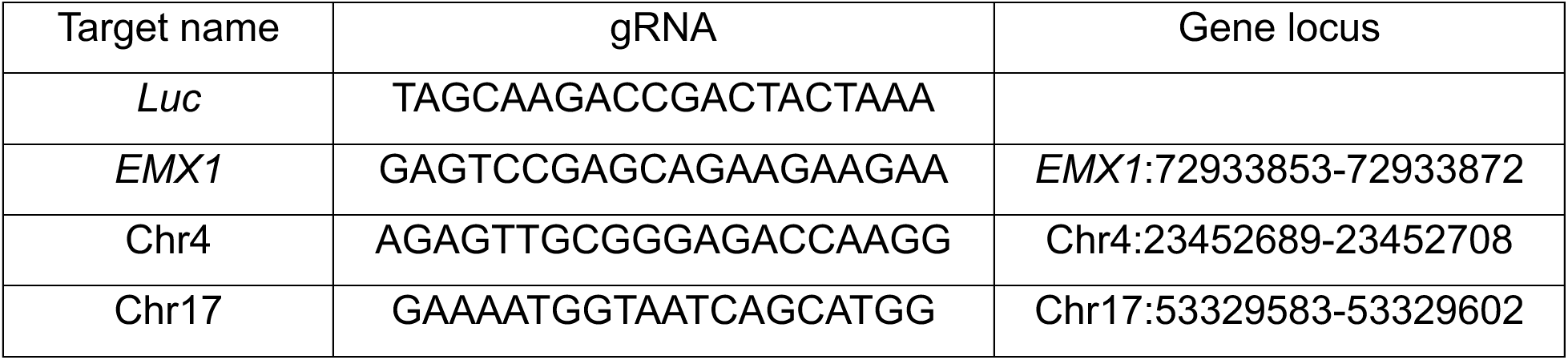
Target sites in this study. All sequences are shown in 5’ to 3’ orientation.

**Supplementary Table 5.**
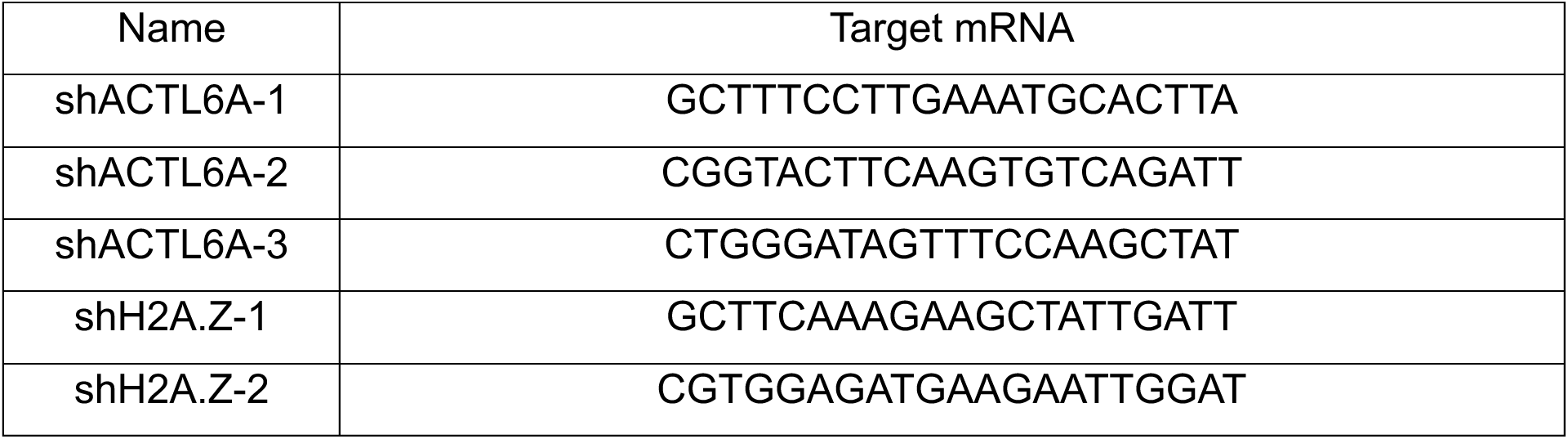
shRNA sequences. All sequences are shown in 5’ to 3’ orientation.

